# Bayesian-estimated hierarchical HMMs enable robust analysis of single-molecule kinetic heterogeneity

**DOI:** 10.1101/404327

**Authors:** Jason Hon, Ruben L. Gonzalez

**Affiliations:** Department of Chemistry, Columbia University, New York, New York 10027, USA

**Keywords:** Single-molecule data analysis, Hierarchical hidden Markov model, Bayesian inference, Variational approximation, Ribosome, Mechanism of translation

## Abstract

Single-molecule kinetic experiments allow the reaction trajectories of individual biomolecules to be directly observed, eliminating the effects of population averaging and providing a powerful approach for elucidating the kinetic mechanisms of biomolecular processes. A major challenge to the analysis and interpretation of these experiments, however, is the kinetic heterogeneity that almost universally complicates the recorded single-molecule signal *versus* time trajectories (*i.e.*, signal trajectories). Such heterogeneity manifests as changes and/or differences in the transition rates that are observed within individual signal trajectories or across a population of signal trajectories. Although characterizing kinetic heterogeneity can provide critical mechanistic information, there are currently no computational methods available that effectively and/or comprehensively enable such analysis. To address this gap, we have developed a computational algorithm and software program, hFRET, that uses the variational approximation for Bayesian inference to estimate the parameters of a hierarchical hidden Markov model, thereby enabling robust identification and characterization of kinetic heterogeneity. Using simulated signal trajectories, we demonstrate the ability of hFRET to accurately and precisely characterize kinetic heterogeneity. In addition, we use hFRET to analyze experimentally recorded signal trajectories reporting on the conformational dynamics of ribosomal pre-translocation (PRE) complexes. The results of our analyses demonstrate that PRE complexes exhibit kinetic heterogeneity, reveal the physical origins of this heterogeneity, and allow us to expand the current model of PRE complex dynamics. The methods described here can be applied to signal trajectories generated using any type of signal and can be easily extended to the analysis of signal trajectories exhibiting more complex kinetic behaviors. Moreover, variations of our approach can be easily developed to integrate kinetic data obtained from different experimental constructs and/or from molecular dynamics simulations of a biomolecule of interest. The hFRET source code, graphical user interface, and user manual can be downloaded as freeware at https://github.com/GonzalezBiophysicsLab/hFRET.

## INTRODUCTION

The kinetic mechanism of a biomolecular process is typically described by specifying the number of states that the biomolecular system samples, the order in which these states are sampled, and the rates of transitions between the sampled states. Over the past twenty years, single-molecule kinetic experiments have emerged as a powerful tool for elucidating such mechanisms^1,2^. This is because the signal *versus* time trajectories (*i.e.*, signal trajectories) that are recorded in such experiments report on the real-time transitions between the states sampled by an individual biomolecule and are therefore free of the population averaging that frequently confounds the analysis of ensemble kinetic experiments. Despite the mechanistically unique and valuable information they provide, single-molecule signal trajectories generally exhibit kinetic heterogeneity, a phenomenon that complicates trajectory analysis and can result in elucidation of incomplete or incorrect kinetic mechanisms^2–4^. Kinetic heterogeneity in a single-molecule kinetic experiment manifests as stochastic, abrupt changes in the rates of transitions observed in individual signal trajectories (*i.e.*, dynamic heterogeneity) and/or as differences in the rates of transitions observed between distinct subpopulations of signal trajectories (*i.e.*, static heterogeneity)^3,5–12^. These effects arise because the signal trajectories recorded in a typical single-molecule kinetic experiment directly detect transitions along only one dimension (*i.e.*, the directly detected dimension) of the complex, multi-dimensional, free-energy landscape that generally governs a biomolecular process^13^. Consequently, transitions along dimensions other than the directly detected dimension (*i.e.*, the indirectly detected dimensions) are projected onto the directly detected dimension, where they materialize indirectly as changes and/or differences in the rates of transitions that are observed in the signal trajectories (Figure 1).

**Figure 1.**
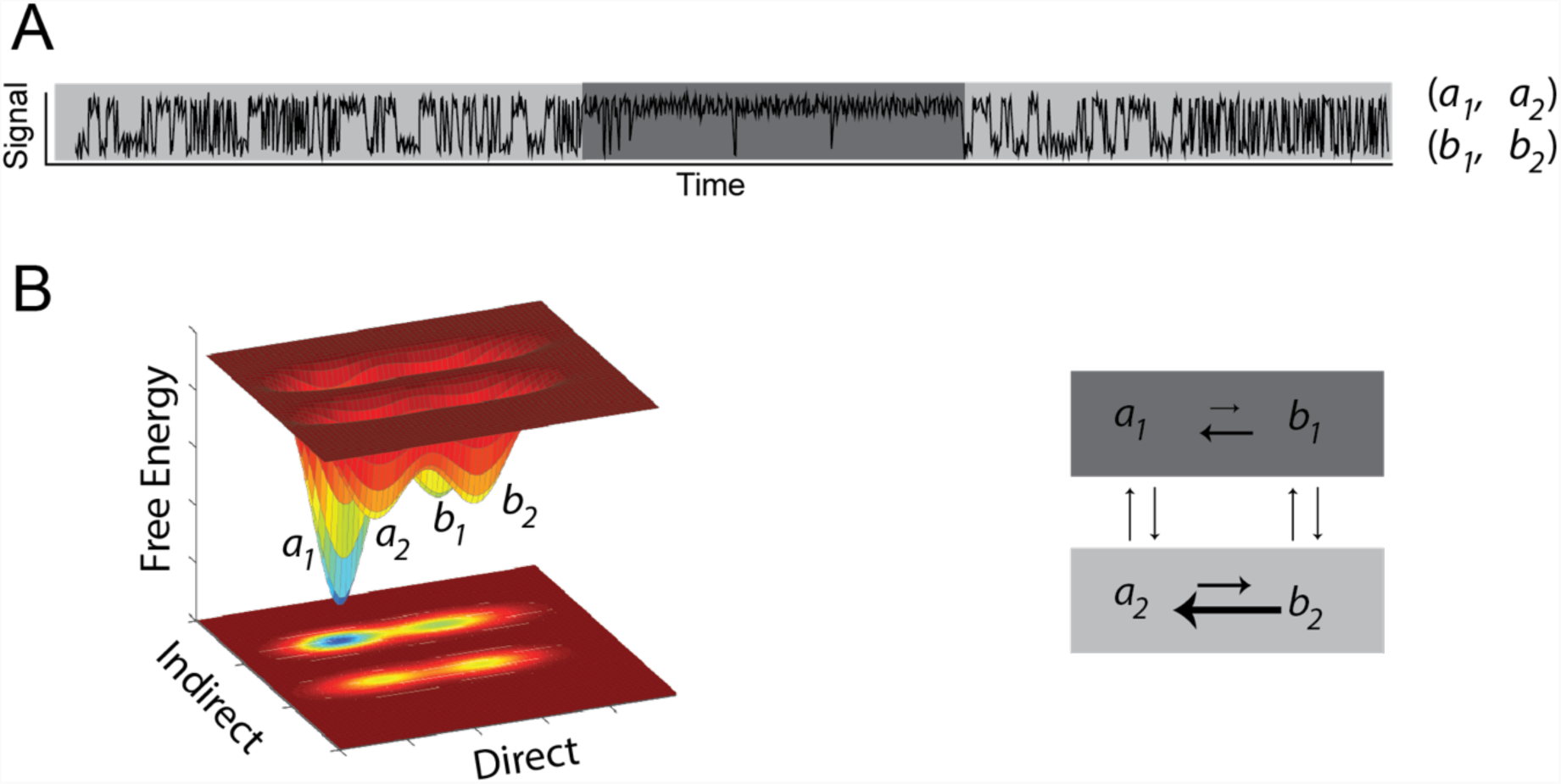
Manifestation and origin of kinetic heterogeneity in single-molecule signal trajectories. (A) A simulated single-molecule signal trajectory composed of contiguous periods, denoted by the variable grayscale backgrounds, in which the rates of transitions between two observable signal amplitudes, denoted as *a*_*n*_ and *b*_*n*_, alternate between two distinct kinetic regimes (where the subscript *n* denotes the signal amplitudes associated with each kinetic regime). (B) The two-dimensional free-energy landscape (left) and corresponding kinetic mechanism (right) used to generate the simulated single-molecule signal trajectory shown in (A). A biomolecule governed by this free-energy landscape can undergo transitions along both a directly observed dimension, or directly detected dimension, denoted by *a*_1_⇄ *b*_1_ and *a*_2_⇄ *b*_2_transitions, and an indirectly observed dimension, or indirectly detected dimension, denoted by *a*_1_⇄ *a*_2_ and *b*_1_⇄ *b*_2_ transitions.

Detecting the presence of transitions along the indirectly detected dimensions of a free-energy landscape and modeling the kinetics of these transitions is of great mechanistic interest. This is because doing so allows identification and characterization of states and subpopulations of a biomolecular system and/or pathways of a biological process that would otherwise be excluded from the kinetic mechanism that is elucidated (*e.g.*, references 5, 6, 12, and 14). Despite its importance, however, detecting and modeling the kinetics of such transitions remains one of the most significant challenges in the analysis and interpretation of single-molecule kinetic experiments^3,5,6,12,14–16^. This is due to inherent limitations^17–23^ in the conventional hidden Markov model (HMM)-based approaches that are widely used to analyze signal trajectories recorded using all of the currently available experimental methods, including single-molecule patch clamp-^24^, fluorescence resonance energy transfer (FRET)-^1,25,26^, force spectroscopy-^1,27^, and field-effect transistor^7,28–31^ experiments. Because the noisy, discretely sampled signal trajectories that are recorded in such experiments^2,32,33^ can be described, to a good approximation, as discrete-time Markov chains^17,34,35^, HMMs have become useful tools for the analysis of these experiments. Nonetheless, it is important to note that the kinetic model employed by an HMM explicitly assumes that the signal trajectory being analyzed only contains transitions that occur along the single, directly detected dimension of a one-dimensional free-energy landscape^36^. This assumption consequently renders HMMs inadequate for the analysis of signal trajectories that additionally contain projections of transitions that occur along the indirectly detected dimensions of a multi-dimensional free-energy landscape. Although several approaches have been developed to circumvent this shortcoming of HMMs in a handful of specific cases^22,37–42^, a solution that can be generally applied to quantify and select between kinetic models of arbitrary complexity has yet to be developed.

To rigorously address this problem, here we have adapted a class of inference tools based on a subclass of Markov chains, known as hierarchical Markov chains, to develop a hierarchical hidden Markov model (HHMM)^43–45^-based approach, which we call hFRET, for the analysis of single-molecule signal trajectories. HHMMs allow signal trajectories to be modeled as though they contain transitions along an arbitrary number of direct and indirectly detected dimensions of a free-energy landscape. Thus, hFRET can be used to identify and characterize kinetic heterogeneity and, correspondingly, to describe biomolecular processes using a hierarchical kinetic mechanism. Moreover, hFRET uses the variational approximation to Bayesian inference^36,46,47^ to estimate the parameters of the HHMMs (*i.e.*, the signal amplitudes of the states and the rates of transitions between states), a method we have previously and successfully used to estimate the parameters of HMMs^20–22^. Because such variational Bayesian methods provide a powerful way to control model complexity^20–22^, hFRET provides a principled approach for selecting the simplest hierarchical kinetic mechanism that best describes the data.

We begin this article by describing the theory underlying hFRET. Using computer-simulated single-molecule signal trajectories derived from a known hierarchical kinetic model and set of parameters, we then assess the accuracy with which hFRET can select the correct hierarchical kinetic model and infer the correct model parameters. Building from the analysis of computer-simulated data derived from a known model, we next use hFRET to analyze experimentally recorded single-molecule FRET (smFRET) data that report on the conformational dynamics of the ribosome, the biomolecular machine that is universally responsible for protein biosynthesis. Our analyses unambiguously reveal the presence of kinetic heterogeneity in single-molecule signal trajectories recorded on several ribosomal complexes and demonstrate that the extent of the heterogeneity depends on the composition of the ribosomal complexes. The approach we present here not only enables researchers to use single-molecule kinetic experiments to develop hierarchical kinetic models describing biological processes of interest, but it also paves the way for the development of closely related approaches that can further expand the data analysis capabilities of the field. Straightforward extensions of the approach presented here, for example, should allow multiple populations of signal trajectories that have been recorded on the same biological process, but using different signals, to be simultaneously analyzed within the context of a single, hierarchical kinetic model. Further extensions should enable the results of single-molecule kinetic experiments on a biological process of interest to be connected to the results of molecular dynamics simulations of the same biological process.

## THEORY

hFRET makes use of hierarchical Markov chains, which are a subclass of Markov chains whose states are parameterized in terms of multiple dimensions as opposed to a single dimension^43^ (Figure 2). The hierarchical Markov chain describing the states and transitions of a biological process, or the “system,” obeys a Kolmogorov-Chapman equation^48^ that propagates a state possessing *D* dimensions 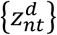 for the n^th^ trajectory at time *t* into the next state in the subsequent time point 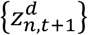, giving rise to the following likelihood function, *L*, for a given population of *N* mutually independent trajectories, each of length *T*_*n*_:

**Figure 2.**
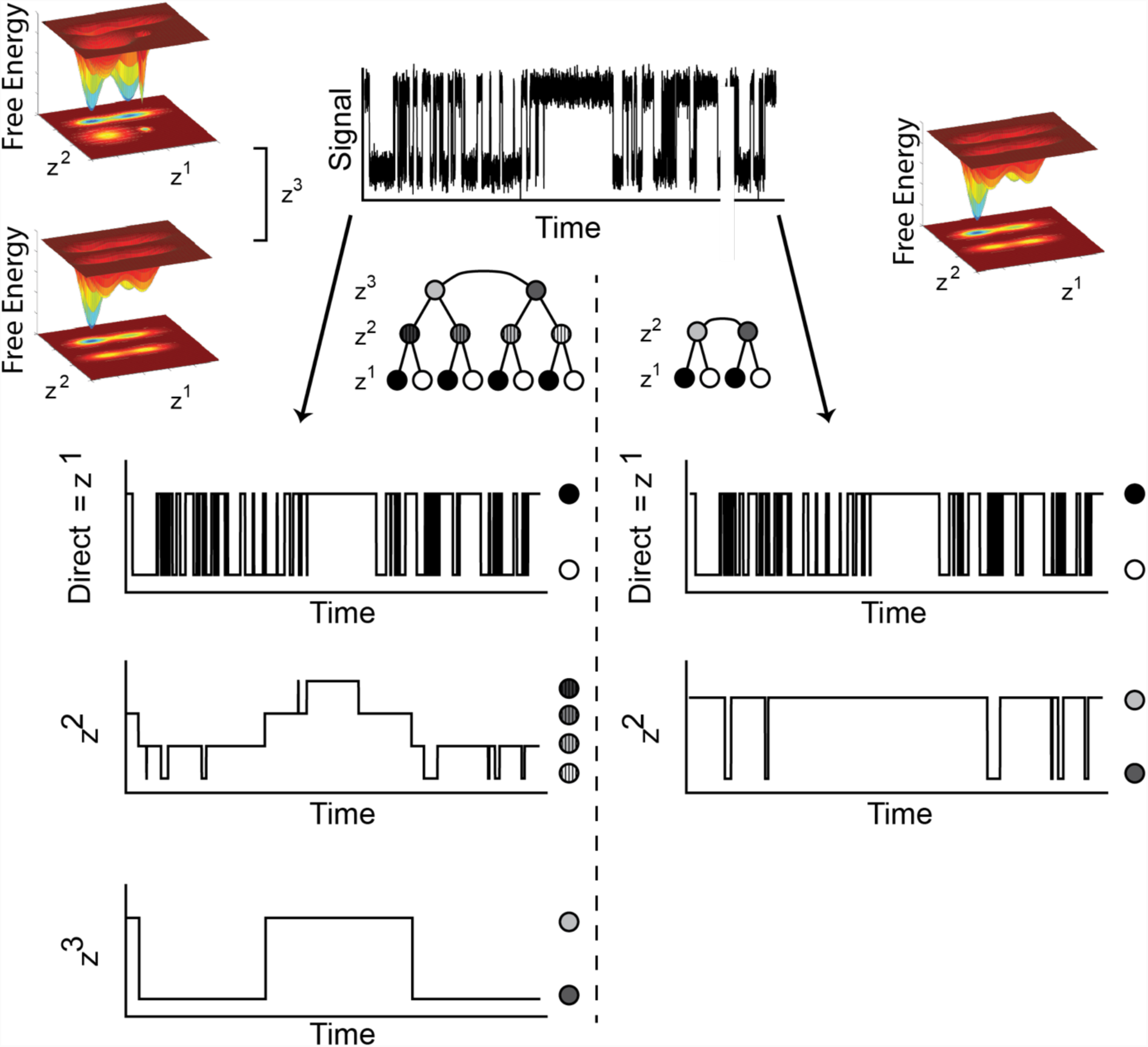
Simulation and analyses of single-molecule signal trajectories generated using a hierarchical Markov chain. A representative, one thousand time-point, single-molecule signal trajectory simulated using a hierarchical Markov chain composed of a directly detected dimension containing two states and two indirectly detected dimensions in which each dimension contains two states (top center). The free-energy landscape and variational Bayesian inference-based analysis generated using the correct HHMM comprised of a directly detected dimension, z^1^, and two indirectly detected dimensions, z^2^ and z^3^ (left) compared to the free-energy landscape and hFRET analysis of a less complex HHMM comprised of a directly detected dimension, z^1^, and only one indirectly detected dimension, z^2^ (right). Both models describe the same transitions along the directly detected dimension but differ in the transitions along and interpretation of indirectly detected dimensions.

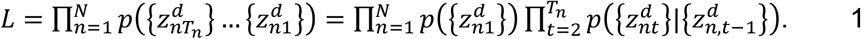

We separate these dimensions into: (i) the directly detected dimension, denoted 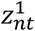 and also referred to as the production level of the state space, which specifies the distribution of observed signal emissions; (ii) the first indirectly detected dimension, denoted 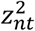, which specifies the distribution of kinetic regimes on the directly detected dimension; and (iii) the arbitrarily higher-order indirectly detected dimensions, denoted 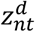, each of which specifies the distribution of kinetic regimes in the indirectly detected dimension that lies directly below it, 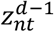. These dimensions are given natural number values that abstractly distinguish the dimensions of a free-energy landscape. The nested, conditional dependencies of this state-space coordinate system may be visualized as a tree of points (see Models 0-5 in Figure 3)^43–45^, which may be thought of as enumerating the order in which the dimensions of a free-energy landscape are specified. Using this convention, the likelihood of the hierarchical Markov chain *L* can be decomposed^44^, beginning with the directly detected dimension and iteratively specifying the abstract values associated with the system on the indirectly detected dimensions:

**Figure 3.**
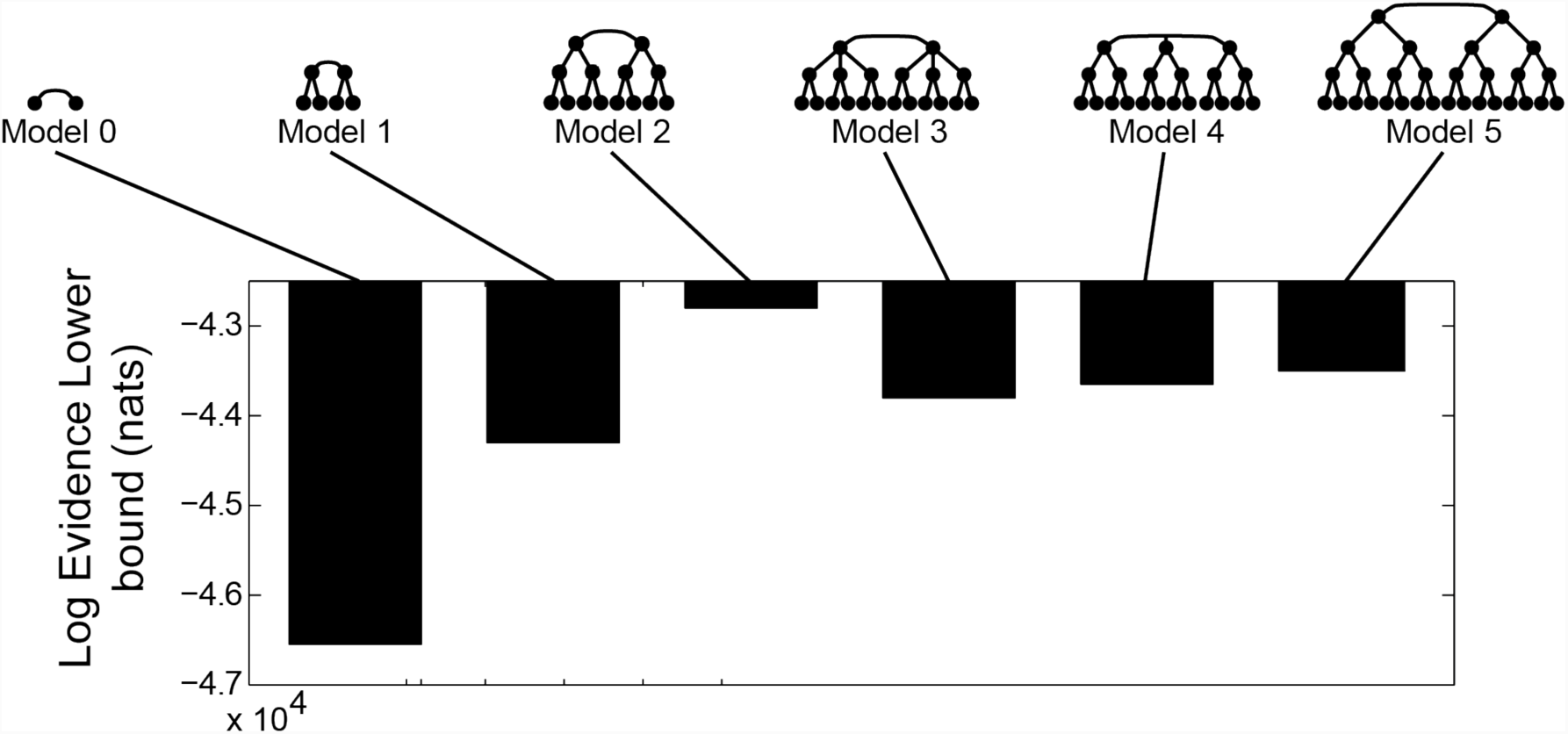
Selection among distinct HHMMs using variational Bayesian inference. Plot of the log evidence lower bound in natural units of information obtained from the variational Bayesian inference-based analysis of six HHMMs, denoted as Models 0-5. Each HHMM is composed of one directly detected dimension containing two states and *n* indirectly detected dimension(s) (where *n* is a number between 0 and 5, as specified by the model numbers denoted along the top of the plot) in which each indirectly detected dimension also contains two states. The tree of points corresponding to each HHMM is depicted along the top of the plot.

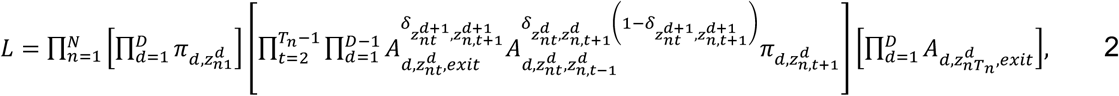

where we have introduced the standard notation:

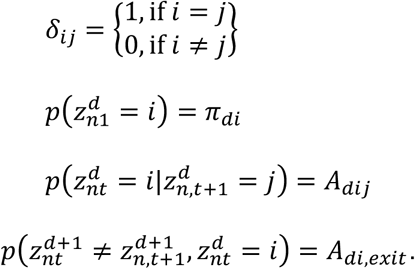

The final statement above represents the probability that the value associated with the system along the indirectly detected dimension *d*, which specifies the distribution of kinetic regimes in the indirectly detected dimension *d* − 1, has transitioned to a new value. We note that a hierarchical kinetic model for static heterogeneity may be derived directly from Equation 2 by simply limiting the equation such that it contains only one indirectly detected dimension and that transitions between kinetic regimes within that indirectly detected dimension are not allowed (presented in greater detail in section S1 of the Supporting Material.)

Equation 2 specifies the hierarchical kinetic model. We use the variational approximation to Bayesian inference^36,46,47^ to specify both the emission distributions and the computational algorithm for estimating the parameters of the HHMM, a procedure that we summarize here and present in greater detail in section S2 of the Supporting Material. Briefly, we seek to maximize the lower bound of the log probability, denoted as the “evidence,” of a set of parameter distributions, denoted as θ, and a set of observations, denoted as {*x*_*nt*_}, given prior information, denoted as *ψ*_0_:

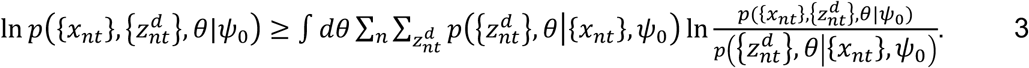

The variational approximation assumes that the coordinates do not depend on the parameter distributions, such that the joint probability may be written as:

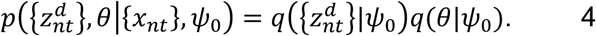

Although here we will assume that the emission distributions are normal distributions, this assumption can be generalized as necessary. Inference of the parameters of an HHMM then proceeds by iteratively locating parameters that optimize a lower bound for the evidence. Iterations proceed by optimizing 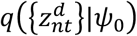, then optimizing *q*(*θ*|*ψ*_0_), and finally calculating the evidence lower bound. Convergence is achieved when the evidence lower bound remains virtually unchanged between iterations.

By utilizing the variational approximation, we can factorize the joint distribution of the kinetic model as follows. First, we simplify the hierarchical Markov chain likelihood in Equation 2 in terms of its transition counts:

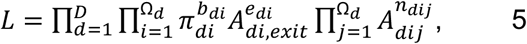

where Ω_*d*_ denotes the number of distinct values of the system along the indirectly detected dimensions in level *d*; *b*_*di*_ denotes the number of transitions resulting in 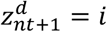 given that 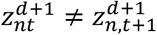 denotes the number of transitions out of 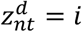 given that 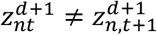; and *n*_*dij*_ denotes the number of transitions from 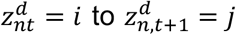. Notably, normalizing *L* implies that the factored distributions over the kinetic parameters decompose into multinomial distributions:

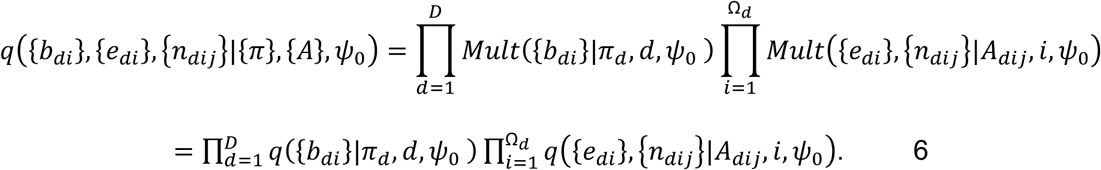

Therefore, considering the state space as a tree of points that are directionally interconnected by conditional relationships, each point can be considered as an independently operating Markov chain, and, to infer the parameters and parameter distributions of the hierarchical kinetic model, it is sufficient to calculate the transition counts specified above. From these parameters, we calculate transition rates between the various free energy minima (see section S3 of the Supporting Material) and therefore quantitatively specify the hierarchical kinetic model.

## RESULTS

### hFRET enables accurate hierarchical kinetic model selection

We first sought to calibrate hFRET by investigating whether it could be used to accurately select among hierarchical kinetic models of systematically increasing complexity. Statistically rigorous model selection is important in data analyses that incorporate indirectly detected dimensions because statistical techniques are the only means for counting and distinguishing amongst alternative models. To test the model selection accuracy of hFRET, we used a specific hierarchical kinetic model to generate a set of simulated signal trajectories, used hFRET to infer the most probable parameters for several models of increasing complexity, and selected the most probable model by calculating and comparing the lower bound of the evidence (see Equation 4). Although the lower bound of the evidence is generally not a rigorous metric for *de novo* model selection^47^, it can be used to compare models as a function of increasing complexity of their state spaces along indirectly detected dimensions. Indeed, when comparing models with an equivalent number of directly detected dimensions but a varying number of indirectly detected dimensions *via* the difference between their evidence lower bound (see section S2 in Supporting Material), the contribution of only the indirectly detected dimensions to the difference between the evidence lower bounds and the evidence *per se* are equivalent^22,36^.

To generate the set of simulated signal trajectories, we used a hierarchical Markov chain composed of one directly detected dimension, two indirectly detected dimensions, and two signal amplitudes, thereby specifying a free-energy landscape consisting of eight distinct free-energy minima, shown in Figure 2 (simulation parameters and sample data provided online at https://github.com/GonzalezBiophysicsLab/hFRET). We subsequently used hFRET to analyze the set of simulated signal trajectories and infer the most probable parameters for six different models (Figure 3). Each of these six models incorporate one directly detected dimension, anywhere from zero to three indirectly detected dimensions, and two signal amplitudes. We have denoted these models as Models 0-5, where the number specifies the number of indirectly detected dimensions that are incorporated by that particular model. We then calculated the lower bound of the evidence for all six models (see section S2 of the Supporting Material.) As expected, the evidence lower bound for the correct model (*i.e.*, Model 2, the model that was used to simulate the data) was significantly larger than that of the incorrect, simpler models (*i.e.*, Models 0-1) as well as the incorrect, more complex models (*i.e.*, Models 3-5). Collectively, the results of these analyses demonstrate that selecting the model with the largest lower bound of the evidence can be used to specify the most parsimonious hierarchical kinetic model that is consistent with the data.

### hFRET allows quantitative characterization of the hierarchical structural dynamics of the ribosome

To demonstrate how hFRET-based analyses of single-molecule kinetic data can be used to characterize the hierarchical dynamics of biomolecular systems, we have used hFRET to analyze signal trajectories from smFRET experiments reporting on the structural dynamics of the bacterial ribosome. The ribosome is the universally conserved, two-subunit, ribonucleoprotein complex that uses aminoacyl-transfer RNA (tRNA) substrates to translate the triplet-nucleotide codon sequence of messenger RNA (mRNA) templates into proteins (Figure 4A). During the elongation stage of translation, addition of each amino acid to the nascent polypeptide chain that is being synthesized by the ribosome proceeds through an elongation cycle comprised of three steps: (i) aminoacyl-tRNA selection, (ii) peptidyl transfer, and (iii) translocation^49^ (Figure 4B).

**Figure 4.**
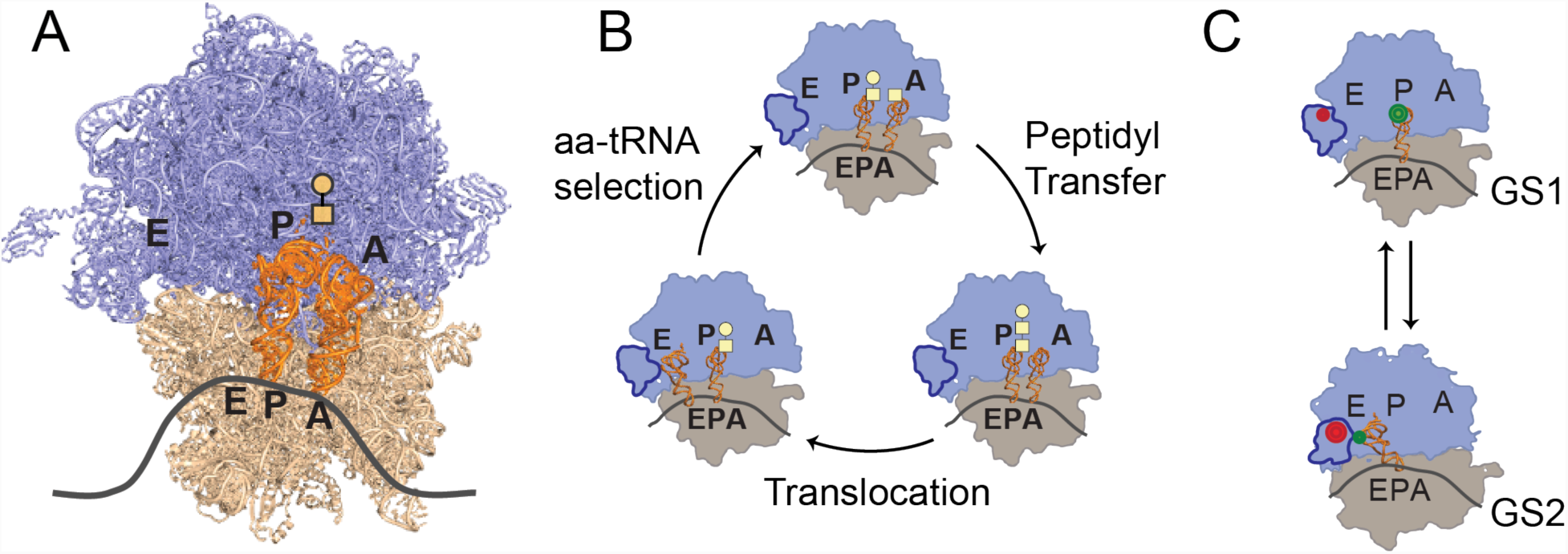
Structure of ribosomal complexes, the translation elongation cycle, and smFRET studies of PRE complexes. (A) X-ray crystallographic structure of an *E. coli* ribosomal complex (PDB ID: 4V51)^62^. The large, or 50S, ribosomal subunit is shown in light blue; the small, or 30S, subunit is shown in tan; the A, P, and E sites on the 50S and 30S subunits are denoted as black letters; the path of the mRNA is denoted as a dark grey curve; the A- and P-site tRNAs are shown in orange; and the location of a hypothetical nascent dipeptide on the A-site tRNA is denoted as yellow shapes. (B) The translation elongation cycle is comprised of aminoacyl-tRNA (aa-tRNA) selection, peptidyl transfer, and translocation. (C) The equilibrium between the GS1 and GS2 conformational states of a PRE complex. The L1-tRNA smFRET signal that is used to report on transitions between GS1 (*E*_FRET_ of ∼0.18) and GS2 (*E*_FRET_ of ∼0.81) in the PRE complex data that are analyzed in the current study is generated using an OH-(Cy3)tRNA^Phe^ within the P site and a (Cy5)L1 within the L1 stalk of the 50S subunit. The Cy3 and Cy5 fluorophores are depicted as green and red circles, respectively.

After undergoing peptidyl transfer, but before undergoing translocation, bacterial ribosomal pre-translocation (PRE) complexes undergo thermally driven, reversible fluctuations between at least two major conformational states that we refer to as global state 1 (GS1) and global state 2 (GS2), establishing a GS1⇄GS2 equilibrium^10^ (Figure 4C). In GS1, the ribosomal subunits occupy their “non-rotated” intersubunit orientation, the L1 stalk element of the 50S subunit occupies its “open” conformation, and the ribosome-bound tRNAs occupy their “classical” configurations. Contrasting with this, in GS2, the ribosomal subunits occupy their “rotated” intersubunit orientation, the L1 stalk element of the 50S subunit occupies its “closed” conformation, and the ribosome-bound tRNAs occupy their “hybrid” configurations.

Previously, we have designed and developed an L1-tRNA smFRET signal by preparing PRE complexes carrying a cyanine (Cy) 3 FRET donor fluorophore-labeled, deacylated, phenylalanine-specific tRNA (OH-(Cy3)tRNA^Phe^) within the ribosomal peptidyl-tRNA binding (P) site and a Cy5 FRET acceptor-labeled ribosomal protein L1 ((Cy5)L1) within the L1 stalk (Figure 4C).^10,50^ Using a total internal reflection fluorescence (TIRF) microscope, we have imaged these PRE complexes at single-molecule resolution, collecting Cy3 and Cy5 fluorescence intensity (*I*_Cy3_ and *I*_Cy5_, respectively) *versus* time trajectories (hereafter referred to as *I*_Cy3_ and *I*_Cy5_ trajectories, respectively) for individual PRE complexes, and using these intensities to calculate the corresponding FRET efficiency (*E*_FRET_) *versus* time trajectory (hereafter referred to as the *E*_FRET_ trajectory), where *E*_FRET_ = *I*_Cy5_ / (*I*_Cy3_+*I*_Cy5_,). The resulting *E*_FRET_ trajectories were observed to spontaneously and stochastically fluctuate between two *E*_FRET_ signal amplitudes, one centered at an *E*_FRET_ of ∼0.18 and the other centered at an *E*_FRET_ of ∼0.81 (see Table 1 for the specific *E*_FRET_ signal amplitudes corresponding to particular PRE complexes). Consistent with the interpretation that PRE complexes undergo thermally driven, reversible fluctuations between GS1 and GS2, the *E*_FRET_ signal amplitudes centered at *E*_FRET_s of ∼0.18 and *∼*0.81 could be assigned to GS1 and GS2, respectively. These assignments were made using structural models of PRE complexes in GS1^51^ and GS2^52^ and the known relationship between *E*_FRET_ and the distance between the donor and acceptor fluorophores *E*_FRET_= 1 / [1 + (R / R_0_)^6^], where R is the distance between the donor and acceptor fluorophores and R_0_, which is also known as the Förster radius, is the distance at which a specific donor and acceptor fluorophore pair exhibit a half-maximal *E*_FRET_ (*i.e.*, an *E*_FRET_ = 0.50).^25^ Further details regarding the design and development of the L1-tRNA smFRET signal and the collection, analysis, and interpretation of L1-tRNA smFRET data can be found in the Materials and Methods section.

**Table 1.** Observed E_FRET_s for each PRE complex. Error bars represent 99% confidence intervals estimated using hFRET.

| PRE Complex | GS1 $E_{\text{FRET}}$ | GS2 $E_{\text{FRET}}$ |
| --- | --- | --- |
| PRE<sup>-A</sup> | 0.168 ± 0.003 | 0.809 ± 0.003 |
| PRE<sup>fMFK</sup> | 0.198 ± 0.002 | 0.818 ± 0.002 |
| PRE<sup>K</sup> | 0.198 ± 0.003 | 0.811 ± 0.003 |

Notably, the L1-tRNA smFRET signal has been used to investigate how the absence, presence, and acylation status of the peptidyl-tRNA in the ribosomal aminoacyl-tRNA binding (A) site modulates the kinetics of GS1→GS2 and GS2→GS1 transitions^10^. Specifically, smFRET data were collected on PRE complexes containing a vacant A site (PRE^−^), carrying an fMet-Phe-Lys-tRNA^Lys^ peptidyl-tRNA in the A site (PRE^fMFK^), and carrying a Lys-tRNA^Lys^ aminoacyl-tRNA in the A site (PRE^K^). Figures 5A, B, and C present plots of representative *I*_Cy3_, *I*_Cy5_, and *E*_FRET_ trajectories corresponding to PRE^−^, PRE^fMFK^, and PRE^K^, respectively. As expected, the *E*_FRET_ trajectories corresponding to all three PRE complexes fluctuate between two *E*_FRET_ signal amplitudes centered at *E*_FRET_s of ∼0.18 and ∼0.81 and corresponding to GS1 and GS2, respectively. hFRET-based analyses of the entire population of *E*_FRET_ trajectories corresponding to either PRE^−^, PRE^fMFK^, or PRE^K^ demonstrated that, for each PRE complex, the most parsimonious hierarchical kinetic model is one in which the directly detected dimension is composed of fluctuations between two directly detected states that are characterized by distinct *E*_FRET_ signal amplitudes centered at *E*_FRET_s of ∼0.18 and ∼0.81 and one indirectly detected dimension is composed of fluctuations between two indirectly detected states that are characterized by distinct rates of transitions between the two signal amplitudes centered at *E*_FRET_s of ∼0.18 and ∼0.81.

**Figure 5.**
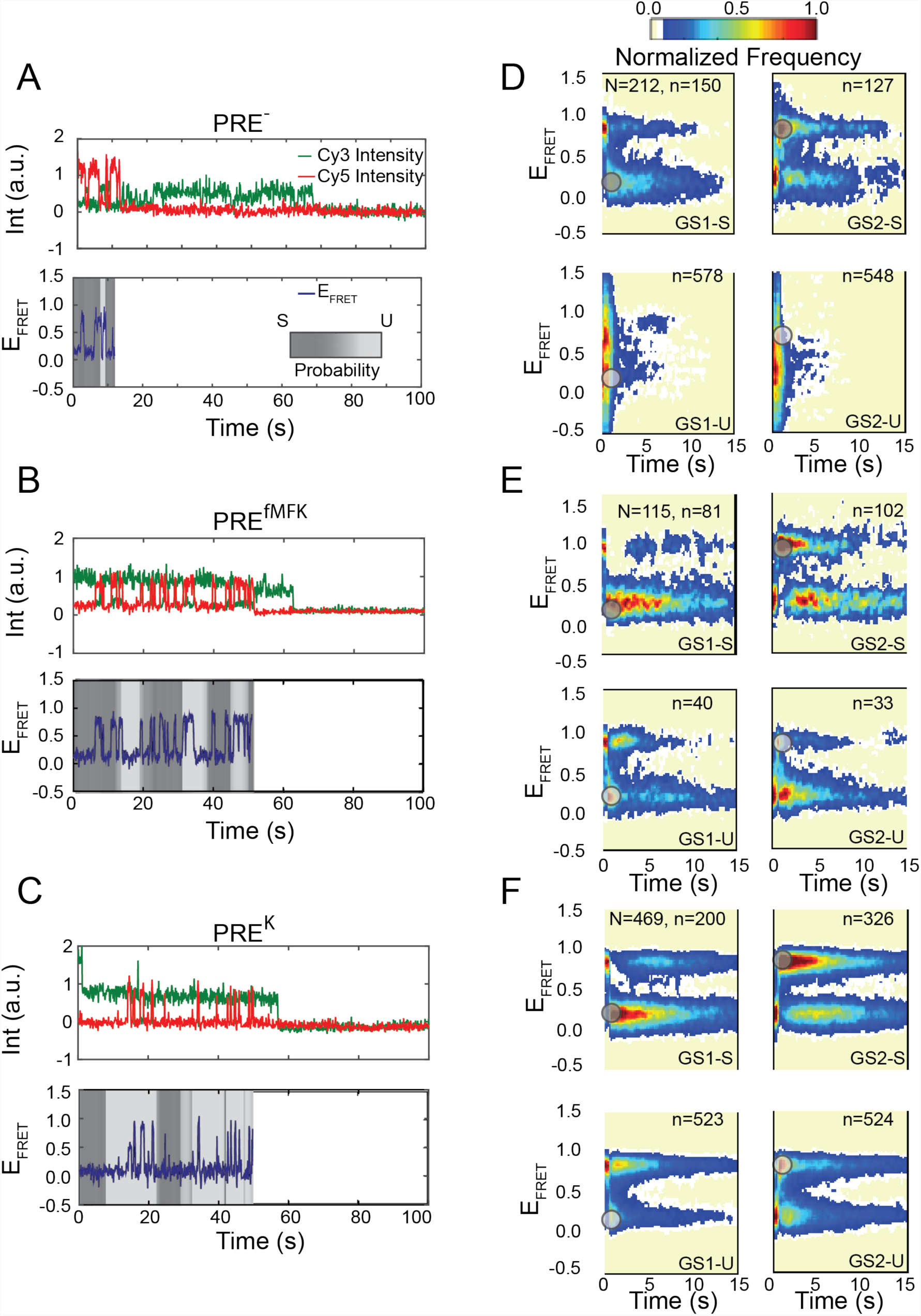
Kinetic heterogeneity of PRE complexes. Representative Cy3- and Cy5 fluorescence intensity trajectories (top) and corresponding *E*_FRET_ trajectories (bottom) are shown for (A) PRE^−^, (B) PRE^fMFK^, and (C) PRE^K^. The backgrounds of the *E*_FRET_ trajectory plots are linearly grayscale-weighted by the probability that a time point belongs to either the S state, dark gray, or the U state, light gray. State recurrence plots (see Results section) for the GS1-S state (upper left-hand plot, initiating at dark gray circle at *E*_FRET_ of ∼0.18), GS2-S state (upper right-hand plot, initiating at dark gray circle at *E*_FRET_ of ∼0.81), GS1-U state (lower left-hand plot, initiating at light gray circle at *E*_FRET_ of ∼0.18), and GS2-U state (lower right-hand plot, initiating at light gray circle at *E*_FRET_ of ∼0.81) of (D) PRE^−^, (E) PRE^fMFK^, and (F) PRE^K^. The ‘N’ denoted at the top of the GS1-S state recurrence plot for each PRE complex specifies the number of distinct *E*_FRET_ trajectories that were used to generate the four state recurrence plots for the corresponding PRE complex. Similarly, the ‘n’ denoted at the top of each state recurrence plot specifies the number of state sub-trajectories that were used to generate the corresponding state recurrence plot.

Consistent with previous work from our group^9,10,50^ and others^53–57^, we interpret the fluctuations between the two directly detected states with distinct *E*_FRET_ signal amplitudes centered at *E*_FRET_s of ∼0.18 and ∼0.81, consistent with the GS1 and GS2 states of the PRE complex, respectively, as reporting on the GS1→GS2 and GS2→GS1 transitions. Moreover, based on a visual inspection of the analyzed *E*_FRET_ trajectories as well as a full kinetic analysis that is presented further below, we interpret the fluctuations between the two indirectly detected states with distinct rates of fluctuations between GS1 and GS2 as reporting on transitions between an indirectly detected state in which excursions to GS1 and GS2 are relatively long-lived and stable, denoted as the Stable (S) state of the PRE complex, and a state in which excursions to GS1 and GS2 are relatively short-lived and unstable, denoted as the Unstable (U) state of the PRE complex.

In order to visually assess the extent to which the individual *E*_FRET_ trajectories recorded for each PRE complex fluctuate between the indirectly detected S and U states (*i.e.*, exhibit dynamic heterogeneity), we generated ‘state recurrence plots’ for the GS1-S, GS2-S, GS1-U, and GS2-U states of each PRE complex (Figures 5D-F). To generate these plots, we first divided each *E*_FRET_ trajectory in the entire population of *E*_FRET_ trajectories corresponding to each PRE complex into GS1-S, GS2-S, GS1-U, and GS2-U sub-trajectories, in which, as a specific example, GS1-S sub-trajectories were defined as those that begin when the *E*_FRET_ signal enters GS1-S and undergo at least one transition to one of the U states (*i.e.*, GS1-U or GS2-U) before returning to GS1-S. GS2-S, GS1-U, and GS2-U sub-trajectories were analogously defined. For each PRE complex, we then plotted a post-synchronized surface contour plot of the time evolution of population FRET for each of the GS1-S, GS2-S, GS1-U, or GS2-U sub-trajectories. As an example, the GS1-S contour plot for each PRE complex was generated by post-synchronizing the GS1-S sub-trajectories from that PRE complex such that the first time point that transitions into GS1-S was assigned to the 0 sec time point on the plot, and then generating a surface contour plot that effectively superimposes all of the post-synchronized transitions into the GS1-S at the 0 sec time point. GS2-S, GS1-U, and GS2-U contour plots were analogously generated. These plots demonstrate that the vast majority of *E*_FRET_ trajectories recorded for each PRE complex fluctuate reversibly between the S and U states and exhibit dynamic heterogeneity. In addition, comparative analyses of these plots suggest that the recurrence times for the U states are particularly sensitive to the presence of an A-site tRNA (compare the two lower plots in Figure 5D to those in Figures 5E and 5F), a qualitative observation that can be more quantitatively characterized using a full kinetic analysis, as described in the next paragraph.

The hierarchical kinetic model described above possesses four distinct free energy minima corresponding to GS1-S, GS2-S, GS1-U, and GS2-U that are connected by eight rate constants (Figure 6). Four of these rate constants connect the states along the directly detected dimension, thus corresponding to the rates of transitions between GS1 and GS2 within the S or U state, denoted as *k*_1,S→2,S,_ *k*_2,S→1,S_, *k*_1,U→2,U_, and *k*_2,U→1,U_ (where the subscripts 1 and 2 denote GS1 and GS2, respectively, and the S and U subscripts denote the S and U states, respectively). The remaining four rate constants connect the states along the indirectly detected dimension, thus corresponding to the rates of transitions between the S and U states within GS1 or GS2, denoted as *k*_1,S→1,U_, *k*_1,U→1,S_, *k*_2,S→2,U_, and *k*_2,U→2,S_. Using this kinetic model, Figure 7 reports the values of these rate constants for the three PRE complexes we have characterized. In all three cases, *k*_1,S→2,S_ was between ∼12- and 54-fold smaller than *k*_1,U→2,U_ and *k*_2,S→1,S_ was between ∼13- and ∼17-fold smaller than *k*_2,U→1,U_, demonstrating that, as qualitatively observed in the individual *E*_FRET_ trajectories (Figure 5), GS1 and GS2 within the S state are significantly more stable than they are within the U state. Moreover, comparison of these rate constants for PRE^−^ and PRE^fMFK^ demonstrates that the PRE complex carrying an fMet-Phe-Lys-tRNA^Lys^ in the A site exhibits a *k*1,S→1,U that is ∼6-fold smaller than that of the PRE complex with an empty A site. Together with slightly smaller, ∼4-fold decreases in *k*_1,U→1,S_, *k*_2,U→1,U_, and *k*_2,S→1,S_, these data demonstrate that the presence of a peptidyl-tRNA in the A site of a PRE complex can modulate rates of transitions along more than one dimension of the multi-dimensional free-energy landscape of a PRE complex. As can be seen by comparing the two lower plots in Figure 5D to those in Figure 5E, these kinetic effects act to increase the recurrence times that are observed for the U states in PRE^fMFK^ *versus* PRE^−^. Similarly, comparison of the rate constants for PRE^K^ and PRE^fMFK^ demonstrates that the PRE complex carrying a Lys-tRNA^Lys^ in the A site exhibits a *k*_2,U→2,S_ that is ∼3-fold larger than that of the PRE complex carrying an fMet-Phe-Lys-tRNA^Lys^ in the A site. Together with a slightly larger, ∼2-fold increase in *k*_1,S→1,U_, these data demonstrate that the acylation status of the A-site tRNA contributes to the ability of the A-site peptidyl-tRNA to modulate the rates of transitions along the indirectly detected dimensions of the free-energy landscape of a PRE complex.

**Figure 6.**
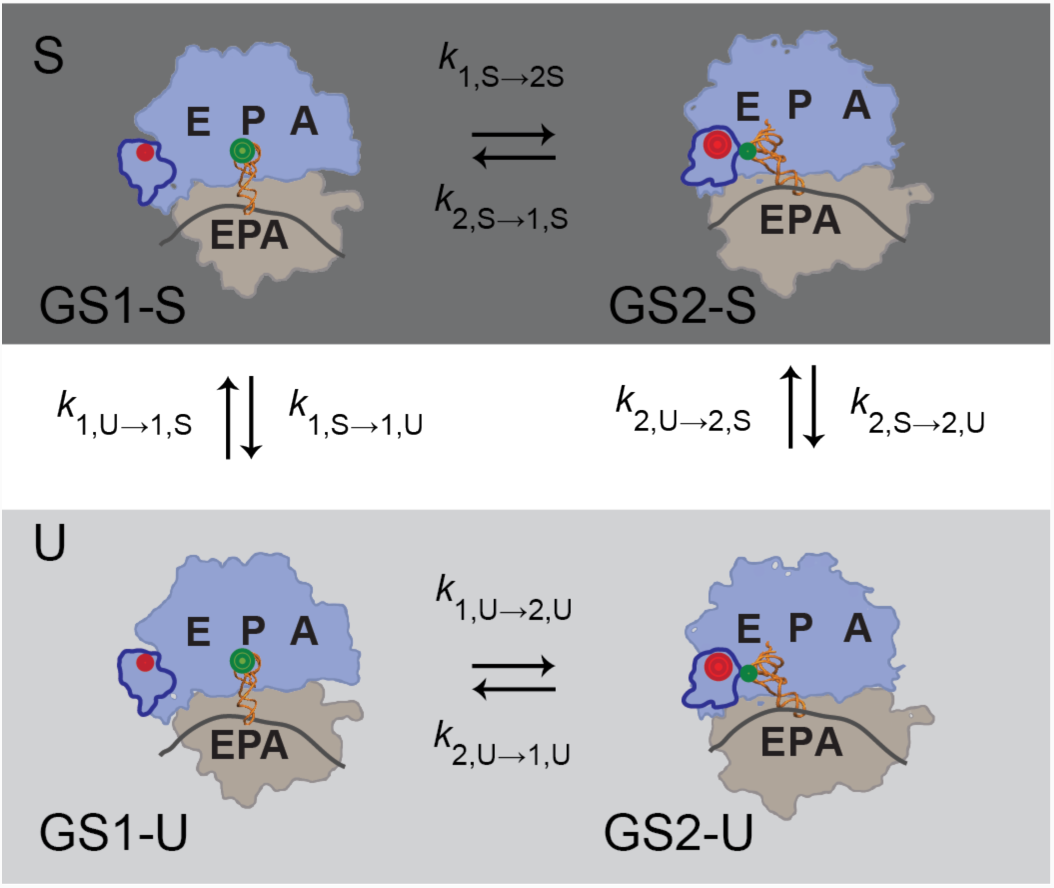
Hierarchical kinetic mechanism of PRE complex dynamics. Hierarchical kinetic mechanism describing the rates of transition between GS1-S, GS2-S, GS1-U, and GS2-U. Transitions between GS1 and GS2 within the S state constant, GS1-S⇄ GS2-S, are enclosed within a box that is shaded in dark gray and transitions between GS1 and GS2 within the U state, GS1-U⇄GS2-U, are enclosed within a box that is shaded in light gray.

**Figure 7.**
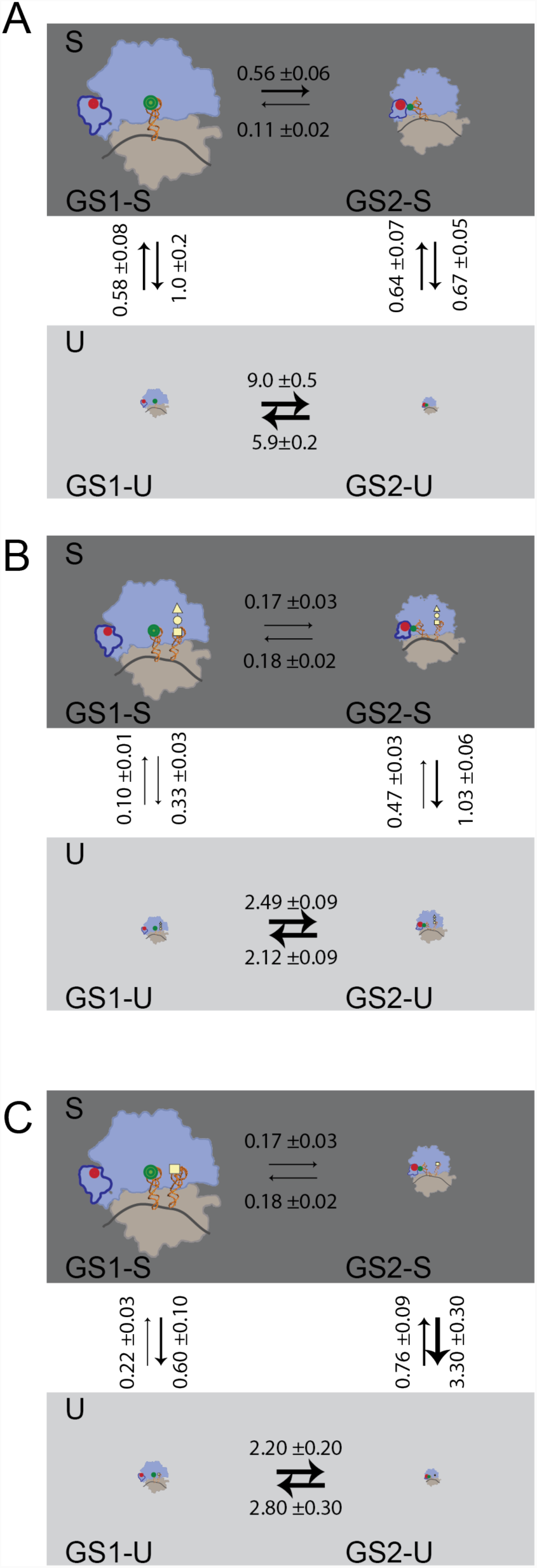
Quantified hierarchical kinetic mechanism of PRE^−^, PRE^fMFK^, and PRE^K^ dynamics. Fully quantified hierarchical kinetic mechanism describing the rates of transitions between GS1-S, GS2-S, GS1-U, and GS2-U for (A) PRE^-^ (B) PRE^fMFK^, and (C) PRE^K^. Note that all rates are in units of s^−1^, error values are 95% confidence intervals, and the relative sizes of the PRE complexes are proportional to the relative occupancies of GS1-S, GS2-S, GS1-U, and GS2-U. The boxes enclosing the various transitions are shaded as in Figure 6.

## DISCUSSION

This work demonstrates a rigorous approach for analyzing single-molecule kinetic data that exhibits kinetic heterogeneity. Specifically, hFRET can be used to identify and kinetically characterize transitions observed in single-molecule signal trajectories whose underlying kinetic model can be best described by a hierarchical Markov chain. This kinetic model describes signal trajectories in which the rates of fluctuations between signal amplitudes that are observed along the dimension of a free-energy landscape that is directly detected in the experiment are modulated by the diffusion of the biomolecule along additional dimensions of the landscape that are not directly detected. hFRET, which uses the variational approximation to optimize the evidence lower bound, enables estimation of the parameters describing a set of hierarchical Markov chains that are consistent with a population of signal trajectories, as well as selection of the hierarchical Markov chain corresponding to the most probable kinetic model that is required to describe the population of signal trajectories. In contrast to existing methods for analyzing single-molecule signal trajectories exhibiting kinetic heterogeneity arising from fluctuations between indirectly detected dimensions of a free-energy landscape^22,37–40^, including a method described as this manuscript was in the final stages of preparation^41^, hFRET enables experimentalists to directly quantify and select between kinetic models generated from hierarchical Markov chains of arbitrary complexity. Thus, hFRET uniquely provides experimentalists with a flexible, comprehensive, and statistically robust method for quantitatively characterizing alternative subpopulations of a biomolecule or biomolecular complex of interest and/or alternative pathways of a biological process of interest^5,6,12,14^.

To demonstrate the ability of hFRET to analyze real-world, single-molecule signal trajectories, we have used it to analyze *E*_FRET_ trajectories reporting on fluctuations between the GS1 and GS2 states of ribosomal PRE complexes. The results of our analyses demonstrate that the most parsimonious hierarchical kinetic model is one that is composed of four states, denoted GS1-S, GS2-S, GS1-U, and GS2-U, in which fluctuations between GS1-S and GS2-S or GS1-U and GS2-U report on transitions along the directly detected dimension of the corresponding free-energy landscape, and fluctuations between GS1-S and GS1-U or GS2-S and GS2-U report on transitions along the indirectly detected dimension of the free-energy landscape. Interestingly, we find that, in all three of the PRE complexes that we investigated, the majority of the *E*_FRET_ trajectories fluctuate reversibly between the S and U states, thereby exhibiting dynamic heterogeneity.

Moreover, our data provide insights into the physical origins of the dynamic heterogeneity of PRE complexes. For example, assuming that the dominant contributions to the energy barriers that separate the S states from the U states in PRE complexes are enthalpic, the fact that *k*_1,S→2,S_ and *k*_2,S→1,S_ are more than one order of magnitude smaller than *k*_1,U→2,U_ and *k*_2,U→1,U_, respectively, for all three PRE complexes, strongly suggests that PRE complexes in the S states are able to form more and/or stronger intramolecular interactions that they can form in the corresponding U states. Given that the rates of transitions between the S and U states, particularly *k*_1,S→1,U_, are sensitive to the presence of a peptidyl-tRNA in the A site of a PRE complex (compare Figure 7A to 7B), we hypothesize that the peptidyl-tRNA at the A site is a major contributor to the differences in the intramolecular interactions that are made in the S states relative to the corresponding U states. This hypothesis is further supported and extended by the observation that the rates of transitions between the S and U states, particularly *k*_2,U→2,S_ and *k*_1,S→1,U_, are sensitive to the acylation status of the tRNA at the A site (compare Figure 6B to 6C). Based on these observations, we propose that the aminoacyl-acceptor stem of the peptidyl-tRNA at the A site of PRE complexes can stochastically and reversibly sample at least two conformations. In one of these conformations, the aminoacyl-acceptor stem makes relatively more and/or stronger interactions with the 50S subunit A and P sites when the peptidyl-tRNA is in its “classical” and “hybrid” configurations, respectively, thereby giving rise to the GS1-S and GS2-S states. In the other conformation, the aminoacyl-acceptor stem makes relatively fewer and/or weaker interactions with the 50S subunit A and P sites when the peptidyl-tRNA is in its “classical” and “hybrid” configurations, respectively, thereby giving rise to the GS1-U and GS2-U states. Thus, our findings strongly suggest that the presence, and likely the identity, post-transcriptional modification status, length and sequence of the covalently attached peptide, *etc.*, of the peptidyl-tRNA at the A site are major contributors to the dynamic heterogeneity of PRE complexes.

Beyond the peptidyl-tRNA at the A site, our results also suggest the existence of additional sources of dynamic heterogeneity in PRE complexes. This is exemplified by the observation that, although it does not carry an A-site tRNA, PRE^−^ also exhibits values of *k*_1,S→2,S_ and *k*_2,S→1,S_ that are more than one order of magnitude smaller than *k*_1,U→2,U_ and *k*_2,U→1,U_, respectively, and likely also forms more and/or stronger intramolecular interactions in the S states than it can form in the corresponding U states. Thus, in analogy with our proposal for the contribution that the peptidyl-tRNA in the A site makes to the kinetic heterogeneity of PRE complexes, it is possible that the aminoacyl-acceptor stem of the deacylated tRNA at the P site of PRE complexes can also stochastically and reversibly sample at least two conformations. Accordingly, such dynamics would modulate the number and/or strength of the interactions that the aminoacyl-acceptor stem of the deacylated tRNA in the P site can make with the 50S subunit P and E sites when the deacylated tRNA is in its “classical” and “hybrid” configurations, respectively, thereby giving rise to the S and U states. Nonetheless, there are alternative possibilities to explain the dynamic heterogeneity we observe in PRE^−^. For example, it is possible that the 30S (and/or 50S) subunit component of an intersubunit interaction can stochastically and reversibly sample at least two conformations that modulate the number and/or strength of the interactions that it could potentially make with its corresponding 50S (and/or 30S) subunit component, thereby giving rise to the S and U states. Thus, in keeping with their compositional and structural complexity, PRE complexes likely contain multiple sources of kinetic heterogeneity and which source dominates the heterogeneity is likely to depend on the precise composition and structure of the particular PRE complex of interest.

Beyond the characterization of static and dynamic heterogeneity in single-molecule signal trajectories, the variational Bayesian HHMM-based approach that underlies hFRET can be adapted to address several other challenges in the field of single-molecule biophysics. Consider, for example, biological processes of greater complexity, in which a sample might exhibit static heterogeneity and each static subpopulation may or may not also exhibit dynamic heterogeneity. Currently, there are no computational methods for analyzing signal trajectories from such samples. Nonetheless, it should be possible to use the variational Bayesian inference-based approach that we have described here together with a set of kinetically non-interconverting hierarchical Markov chains to develop an hFRET-like algorithm that can be used to analyze signal trajectories of such complexity. In a second example, we note that there are currently no standard computational methods available for elucidating the single, most parsimonious kinetic model that is fully consistent with the signal trajectories from multiple datasets in which the signal trajectories from each dataset report on the conformational dynamics of a distinct structural element of a biomolecule or biomolecular complex of interest (*e.g*., the various smFRET datasets associated with the different FRET donor and acceptor pairs that have been developed and used to investigate the intersubunit-, L1 stalk-, and tRNA dynamics of PRE complexes^13^). Using a variational Bayesian HHMM-based approach that builds on hFRET so as to treat the signal trajectories from each dataset as reporting on a different dimension of the same free-energy landscape should allow the single, most parsimonious kinetic model that is fully consistent with all of the datasets to be selected. A final example is really an extension of the second example, in which the experimental signal trajectories from one dataset are replaced by signal trajectories derived from molecular dynamics (MD) simulations. Such an approach would provide a robust method for integrating experimental and MD simulation data into a single kinetic model. Although the sampling period of the detectors that are used in most single-molecule biophysical experiments (∼1-100 milliseconds sampled for ∼1-100 seconds) is currently much longer than the maximum time of an MD simulation (∼10 milliseconds^58^), rapid improvements in the detectors and increases in computational power will soon allow the timescales accessible to single-molecule experiments and MD simulations to be bridged.

## MATERIALS AND METHODS

### Generation of simulated signal trajectories using a specific hierarchical kinetic model

One thousand signal trajectories composed of one thousand time points each were simulated by randomly drawing each signal trajectory from a hierarchical Markov chain. This hierarchical Markov chain was composed of a directly detected dimension containing two states characterized by two distinct signal amplitudes and two indirectly detected dimensions in which each dimension contained two distinct states (*i.e.*, the hierarchical Markov chain shown in the left-hand side of Figure 2). Gaussian-distributed noise to a final signal-to-noise ratio of 5:1 was then added to each of the simulated signal trajectories. The source code for generating simulated signal trajectories, as well as sample simulated signal trajectories, can be found together with the hFRET source code, graphical user interface, and user manual at https://github.com/GonzalezBiophysicsLab/hFRET.

### Collection and analysis of smFRET data

The L1-tRNA smFRET data that was analyzed using hFRET in the current study consists of datasets that had been previously collected; analyzed using a different, maximum likelihood-estimated HMM-approach^18^; and interpreted and reported by Fei *et al*^10^. Briefly, the L1-tRNA smFRET signal was generated by preparing PRE complexes carrying an OH-(Cy3)tRNA^Phe^ within the P site and (Cy5)L1 within the L1 stalk of the 50S subunit. OH-(Cy3)tRNA^Phe^ was prepared by site-specifically labeling the dihydrouridine at position 47 of OH-tRNA^Phe^ (Sigma) with Cy3. (Cy5)L1 was prepared by site-specifically labeling an introduced cysteine at position 202 in a recombinantly overexpressed and purified single-cysteine variant of *Escherichia coli* ribosomal protein L1 with Cy5. (Cy5)L1-labeled 50 subunits were subsequently generated by reconstituting (Cy5)L1 into 50S subunits that had been purified from an *E. coli* strain lacking the gene encoding ribosomal protein L1.

As discussed in more detail elsewhere^10^, three PRE complexes were assembled onto mRNAs containing a biotin moiety at the 5’ terminus. PRE^−^, which carried an OH-(Cy3)tRNA^Phe^ at the P site and a vacant A site, was prepared by delivering puromycin, a ribosome-targeting inhibitor of protein synthesis, to the A site of a ribosomal elongation complex carrying an fMet-Phe-tRNA^Phe^ at the P site and a vacant A site. Puromycin is an analog of the 3’-terminal residue of aminoacyl-tRNA that binds to the A site of the peptidyl transferase center of the 50S subunit, acts as an acceptor substrate in the peptidyl transfer reaction with the fMet-Phe-(Cy3)tRNA^Phe^ at the P site, and rapidly dissociates from the 50S subunit, thereby generating a PRE complex containing a deacylated OH-(Cy3)tRNA^Phe^ in the P site and a vacant A site (*i.e.*, PRE^−^)^59^. PRE^fMFK^, which carried an OH-(Cy3)tRNA^Phe^ at the P site and an fMet-Phe-Lys-tRNA^Lys^ at the A site, was prepared by delivering a ternary complex composed of elongation factor (EF) Tu, GTP, and Lys-tRNA^Lys^ (EF-Tu(GTP)Lys-tRNA^Lys^) to the A site of a ribosomal elongation complex carrying an fMet-Phe-(Cy3)tRNA^Phe^ at the P site and a vacant A site. Once accommodated into the A site, Lys-tRNA^Lys^ acts as an acceptor substrate in the peptidyl transfer reaction, thereby generating a PRE complex carrying an OH-(Cy3)tRNA^Phe^ at the P site and an fMet-Phe-Lys-tRNA^Lys^ at the A site (*i.e.*, PRE^fMFK^). PRE^K^, which carried an an OH-(Cy3)tRNA^Phe^ at the P site and an Lys-tRNA^Lys^ at the A site, was prepared by delivering EF-Tu(GTP)Lys-tRNA^Lys^ to the A site of a PRE complex identical to PRE^−^ and thereby carrying an OH-(Cy3)tRNA^Phe^ at the P site and a vacant A site PRE^−^. Although it accommodates into the A site, the lack of a peptidyl moiety on the OH-(Cy3)tRNA^Phe^ at the P site prevents Lys-tRNA^Lys^ from acting as an acceptor substrate in the peptidyl transfer reaction, thereby generating a PRE complex carrying an OH-(Cy3)tRNA^Phe^ at the P site and a Lys-tRNA^Lys^ at the A site (*i.e.*, PRE^K^).

As previously described in greater detail^10,60^, a laboratory-built, prism-based, wide-field single-molecule total internal reflection fluorescence (TIRF) microscope was used to image the three PRE complexes in a Tris-Polymix imaging buffer composed of 50 mM tris(hydroxymethyl)aminomethane acetate, 100 mM potassium chloride, 5 mM ammonium acetate, 0.5 mM calcium acetate, 15 mM magnesium acetate, 6 mM β-mercaptoethanol, 5 mM putrescine dihydrochloride, and 1 mM spermidine free base at a pH_25°C_ of 7.5 that was supplemented with an oxygen-scavenging system (1% β-D-glucose, 25 units/mL glucose oxidase, and 250 units/ml catalase)^10^. Briefly, each PRE complex was tethered to the surface of a microfluidic TIRF microscopy observation flowcell that had been passivated with a mixture of polyethylene glycol (PEG) and biotinylated PEG and had been treated with streptavidin. Cy3 fluorophores were directly excited using a 532-nm laser excitation source (CrystaLaser) and fluorescence emissions from both Cy3 and Cy5 were collected using a 1.2 numerical aperture/60× objective (Nikon), wavelength separated using a two-channel imaging system (Photometrics Inc.), and recorded as a movie using a back-illuminated, electron-multiplying, charge-coupled device camera (Photometrics) operating at an acquisition time of 50 ms per frame.

As detailed in our previous report^10^, individual pairs of Cy3- and Cy5 fluorescence intensity *versus* time trajectories (*i.e.*, intensity trajectories) reporting on the conformational dynamics of single PRE complexes were generated using laboratory-written, MATLAB-based, image-analysis software. Each pair of Cy3- and Cy5 fluorescence intensity trajectories was: (i) truncated at the first time point at which either fluorophore ‘photobleached’ (*i.e.*, underwent an apparently irreversible loss of fluorescence intensity), (ii) baseline corrected by subtracting the average intensity of the last ten time points following the photobleaching event of either the Cy3 or Cy5 fluorophore, and (iii) spectral cross-talk corrected by decreasing the Cy5 fluorescence intensity at each time point by 7% of the Cy3 fluorescence to account for the 7% bleed-through of Cy3 fluorescence emission into the Cy5 channel. The photobleaching-, baseline-, and spectral cross-talk corrected pairs of Cy3- and Cy5 fluorescence intensity trajectories were then used to generate individual *E*_FRET_ trajectories by using the Cy3 intensity at each timepoint, *I*_*cy3*_ (*t*), and the Cy5 intensity at each timepoint, *I*_*cy5*_ (*t*), to calculate the *E*_FRET_ at each time point as *E*_FRET_(*t*) = *I*_*cy5*_(*t*)/(*I*_*cy5*_(*t*) + *I*_*cy3*_(*t*)). Assuming the quantum yield of the Cy3 fluorophore and the rotational motion of the Cy3- and/or Cy5 fluorophores are constant throughout the experiment, changes in *E*_FRET_ are proportional to changes in the distance between the Cy3 and Cy5 fluorophores as given by *E*_FRET_(*R*) = 1/(1 + (*R*/*R*_0_)^>6^), where *R* is the distance between the Cy3 and Cy5 fluorophores and *R*_0_ is the Forster radius, which is ∼54 Å under our conditions^25,61^. The resulting *E*_FRET_ trajectories were analyzed using hFRET and visualized using state recurrence plots as discussed in the Results section. The hFRET source code, graphical user interface, and user manual can be found together with source code for generating simulated signal trajectories and sample simulated signal trajectories at https://github.com/GonzalezBiophysicsLab/hFRET.

## SUPPORTING INFORMATION

A detailed model, update equations, and description of the computational algorithm is available in the Supporting Information. Code and sample data are available at https://github.com/GonzalezBiophysicsLab/hFRET.

## AUTHOR CONTRIBUTIONS

J.H. and R.L.G. designed the research, analyzed the data, and wrote the paper.

## ACKNOWLEDGEMENTS

The authors thank Dr. Kelvin Caban and Dr. Colin Kinz-Thompson for their comments on the manuscript. This work was supported by a National Science Foundation CAREER award (MCB 0644262), NIH grants under award numbers GM107417, GM084288, GM119386; and an American Cancer Society grant under award number RSG-09-053-01-GMC to RLG.

## COMPETING FINANCIAL INTERESTS

The authors declare no conflicts of interest.

